# Zika virus infection of pregnant *Ifnar1*^−/−^ mice triggers strain-specific differences in fetal outcomes

**DOI:** 10.1101/2021.05.14.444269

**Authors:** Ellie K. Bohm, Jennifer T. Vangorder-Braid, Anna S. Jaeger, Ryan V. Moriarty, John J. Baczenas, Natalie C. Bennett, Shelby L. O’Connor, Michael K. Fritsch, Nicole A. Fuhler, Kevin K. Noguchi, Matthew T. Aliota

## Abstract

Zika virus (ZIKV) is a flavivirus that causes a constellation of adverse fetal outcomes collectively termed Congenital Zika Syndrome (CZS). However, not all pregnancies exposed to ZIKV result in an infant with apparent defects. During the 2015-2016 American outbreak of ZIKV, CZS rates varied by geographic location. The underlying mechanisms responsible for this heterogeneity in outcomes have not been well defined. Therefore, we sought to characterize and compare the pathogenic potential of multiple Asian/American-lineage ZIKV strains in an established *Ifnar1^−/−^* pregnant mouse model. Here, we show significant differences in the rate of fetal demise following maternal inoculation with ZIKV strains from Puerto Rico, Panama, Mexico, Brazil, and Cambodia. Rates of fetal demise broadly correlated with maternal viremia but were independent of fetus and placenta virus titer, indicating that additional underlying factors contribute to fetus outcome. Our results, in concert with those from other studies, suggest that subtle differences in ZIKV strains may have important phenotypic impacts. With ZIKV now endemic in the Americas, greater emphasis needs to be placed on elucidating and understanding the underlying mechanisms that contribute to fetal outcome.

**IMPORTANCE:** Zika virus (ZIKV) actively circulates in 89 countries and territories around the globe. ZIKV infection during pregnancy is associated with adverse fetal outcomes including birth defects, microcephaly, neurological complications, and even spontaneous abortion. Rates of adverse fetal outcomes vary between regions, and not every pregnancy exposed to ZIKV results in birth defects. Not much is known about how or if the infecting ZIKV strain is linked to fetal outcomes. Our research provides evidence of phenotypic heterogeneity between Asian/American-lineage ZIKV strains and provides insight into the underlying causes of adverse fetal outcomes. Understanding ZIKV strain-dependent pathogenic potential during pregnancy and elucidating underlying causes of diverse clinical sequelae observed during human infections is critical to understanding ZIKV on a global scale.

## INTRODUCTION

Zika virus (ZIKV) exposure during pregnancy can cause a constellation of adverse fetal outcomes, collectively termed congenital Zika syndrome (CZS). However, a substantial proportion of pregnancies with in-utero ZIKV exposure result in babies without apparent defects. Only an estimated 5-15% of infants have ZIKV-related birth defects (1–3). Importantly, infants who are born apparently healthy can manifest developmental and neurocognitive deficits months to years after birth (4–7), even if maternal exposure resulted in asymptomatic infection (8). Furthermore, there was an unequal distribution of ZIKV cases and severe outcomes in all areas where ZIKV emerged in the Americas, demonstrating that risk of CZS varied over time and with geographic location (reviewed in (9)). For example, the rate of microcephaly differed between French Polynesia (1%) (10), the U.S. Territories and Freely Associated States (5-6%) (11), and the Dominican Republic (11%) (7). Within Brazil, the rate of microcephaly varied between São Paulo (0%) (12), Pernambuco (2.9%) (13), Rio de Janeiro (3.5%) (14), Southeast Brazil (1.5%) and Northeast Brazil (13%) (15). However, it should be noted that accurate diagnosis of microcephaly requires multiple measures after birth and the use of inconsistent definitions of cases and complications can bias reporting (9). For example, initial microcephaly rates were overestimated in Brazil before INTERGROWTH-21st reference-based standards were implemented (16).

Microcephaly is not the only adverse birth outcome that results from gestational ZIKV infection (17), and these rates varied as well. The U.S. Territories and Freely Associated States reported birth defects in 14% of ZIKV-exposed pregnancies (11). Pernambuco, Brazil reported adverse outcomes in 20% of exposed pregnancies (13), whereas São Paulo reported a 28% rate of adverse neurological outcomes (12). Strikingly, in Rio de Janeiro, 42% of infants born to ZIKV-exposed pregnancies had adverse outcomes; however, this study used a broader definition for ZIKV-associated outcomes (14). Because current diagnostic testing remains suboptimal and inconsistent for the detection of congenital ZIKV infection (18), the relative risk of CZS in infants from ZIKV-exposed pregnancies remains unknown, and it remains unknown whether the risk is equal in different geographic areas. Was the unequal distribution in CZS incidence over time and region stochastic or were there other factors that influenced these regional differences? A provocative explanation for the appearance of CZS in the Americas is that contemporary ZIKV strains evolved from strains that cause fetal lethality to those that cause birth defects and this may have facilitated recognition of ZIKV’s ability to harm the developing fetus (19). Whether ongoing virus evolution during geographic spread in the Americas gave rise to phenotypic variants that differ in their capacity to harm the developing fetus remains an open question.

Large case-control studies of pregnant women may prove useful for determining whether infecting ZIKV genotype affects overall pathogenesis during pregnancy. However, these types of studies are observational and are complicated by participant heterogeneity, including history of infection with other flaviviruses, and the precise time, dose, and genetic makeup of the infecting virus. We therefore aimed to better understand heterogeneity in ZIKV-associated pregnancy outcomes by investigating whether there are Asian/American-lineage strain-specific phenotypic differences by using mice lacking type I interferon signaling (*Ifnar1^−/−^*). Although there are limitations regarding the translational relevance of this model, transplacental ZIKV infection and fetal damage have been demonstrated (20–23), and it has been used to compare maternal infection parameters, placental pathology, fetal infection, and outcomes between ZIKV strains and the closely-related Spondweni virus (20–22). Congenital ZIKV studies in pregnant mouse models have used a variety of virus strains, as well as timing, route, and dose of inoculation (22–25). This heterogeneity in design has made it difficult to compare results across mouse studies because both inoculation dose and time of ZIKV exposure during pregnancy play a role in determining fetal outcomes (26). Therefore, we assessed fetal outcomes following infection by a panel of five geographically distinct, low-passage Asian/American-lineage ZIKV strains at embryonic day 7.5 (E7.5). Here, we found that all ZIKV strains infected the placenta but varied in their capacity to cause overt fetal harm, suggesting that there is phenotypic heterogeneity in pregnancy outcomes that is dependent on the infecting ZIKV genotype.

## RESULTS

### ZIKV replication kinetics in maternal serum are broadly similar among Asian/American-lineage strains

To perform a comprehensive phenotypic characterization of ZIKV infection during pregnancy, we assembled a set of five recently isolated, low-passage ZIKV strains based on their geographic distribution in the Americas and minimal passage history. Our ZIKV panel included four epidemic strains from the American-sublineage (Puerto Rico-2015, PR; Panama-2015, PAN; Mexico-2016, MEX; and Brazil-2015, BRA) and a non-epidemic strain from the Asian-lineage (Cambodia-2010, CAM). ZIKV-PR, ZIKV-PAN, ZIKV-MEX, and ZIKV-BRA share >99.5% genome-wide nucleotide identity, and ZIKV-CAM shares over 98% genome-wide nucleotide identity, resulting in only 4-18 amino acid differences between strains (**Tables 1** **and** **2**).

**Table 1:** Total number of amino acid differences between strains and (percent difference in amino acid identity).

|  | CAM | BRA | MEX | PAN |
| --- | --- | --- | --- | --- |
| PR | 17 (0.50%) | 4 (0.12%) | 9 (0.26%) | 7 (0.20%) |
| PAN | 16 (0.47%) | 5 (0.15%) | 8 (0.23%) |  |
| MEX | 18 (0.53%) | 7 (0.20%) |  |  |
| BRA | 15 (0.44%) |  |  |  |

**Table 2:**
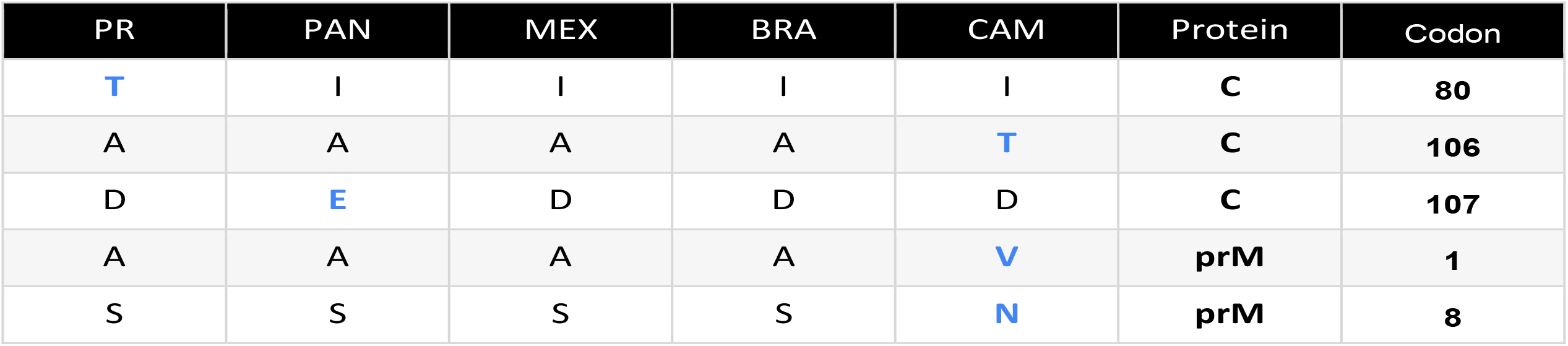
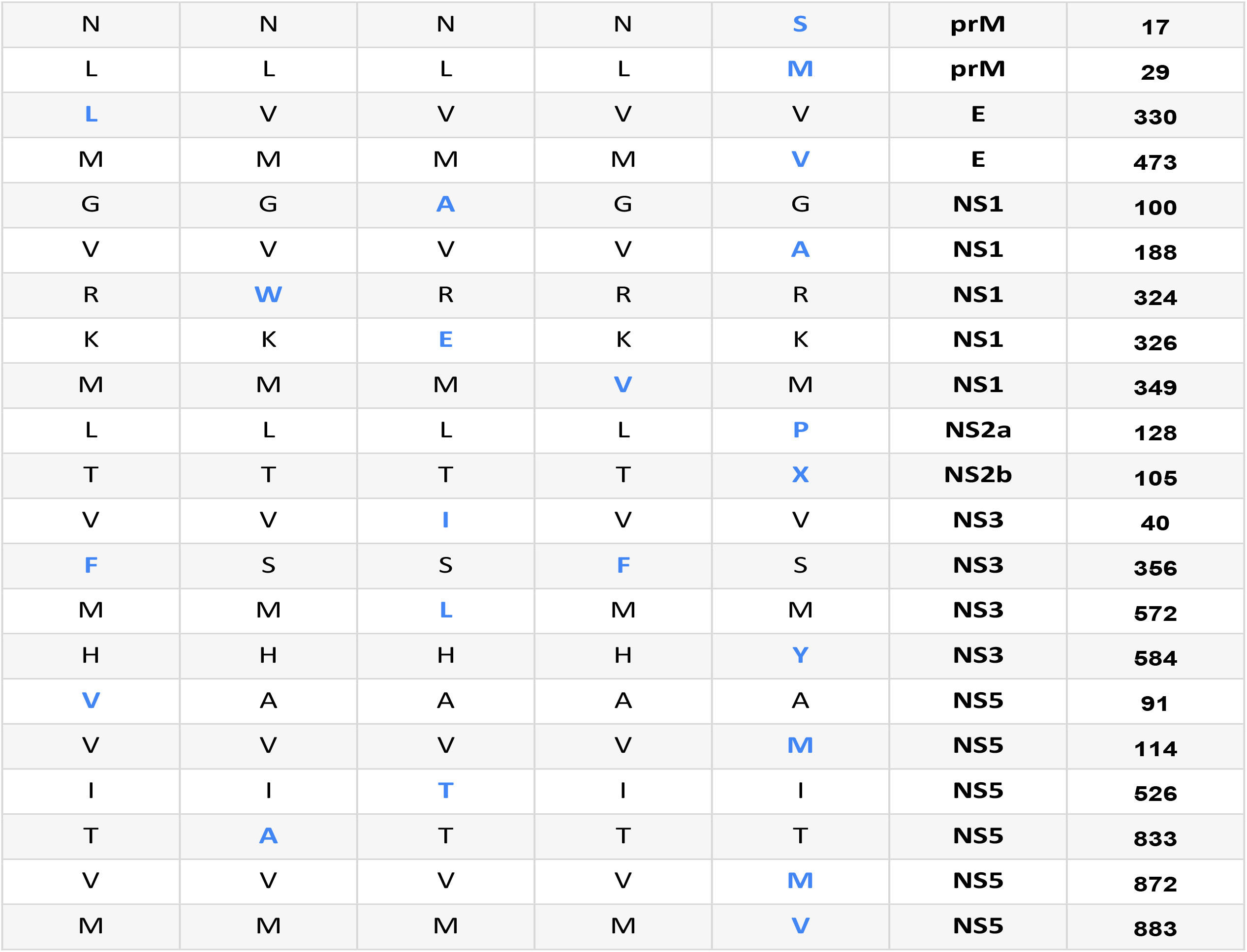
Differences in amino acid sequences across Asian/American-Lineage ZIKV strains. PR (PRVABC59; GenBank:AMC13911.1), PAN (259249; GenBank:ANB66183), MEX (R116265; GenBank:AOG18296.1), CAM (FSS13025; GenBank:AFD30972), BRA (Paraiba_01; GenBank:ANH10698.1).

To characterize the range of pathogenic outcomes and assess the effect of ZIKV strain on in-utero exposure, we utilized a well-established murine pregnancy model for ZIKV (20, 21). *Ifnar1*^−/−^ dams were time-mated with wildtype (WT) males to produce fetuses and a maternal-fetal interface (MFI) with intact interferon (IFN) signaling. Pregnant *Ifnar1*^−/−^ dams were inoculated with 1×10^3^ PFU of ZIKV-PR, ZIKV-PAN, ZIKV-MEX, ZIKV-BRA, or ZIKV-CAM via subcutaneous footpad inoculation at embryonic day 7.5 (E7.5), corresponding to the mid-to-late first trimester in humans (27). Based on results from our past studies (20, 21), we chose this dose to minimize the potential confounding impacts of maternal illness on fetal outcomes. Maternal serum samples were collected at 2, 4, and 7 days post inoculation (dpi) to confirm infection and examine viremia kinetics (**Figure 1A**). All dams were productively infected with detectable viremia by 4 dpi for all groups. ZIKV-CAM replicated to significantly higher titers at 2 dpi compared to ZIKV-PAN and ZIKV-MEX (One-way ANOVA with Tukey’s multiple comparisons, *p* = 0.0199 and *p* = 0.0392), whereas maternal viremia was significantly lower in ZIKV-BRA-inoculated dams compared to all other treatment groups at this timepoint. By 4 dpi, ZIKV-BRA had replicated to significantly higher titers compared to ZIKV-PAN and ZIKV-MEX (*p* = 0.0026 and *p* <0.0001). Maternal viremia in the ZIKV-CAM group also was significantly higher compared to the ZIKV-MEX group at 4 dpi (*p* = 0.0024). Overall, maternal viremia reached similar levels before being cleared to the limit of detection by 7 dpi in all groups. Due to the impact of COVID-19, ZIKV-PAN maternal serum samples at 7 dpi were not collected. Dams were monitored daily for clinical signs until the time of necropsy at E14.5 (7 dpi) and no overt clinical signs were observed in any virus- or PBS- inoculated dams.

**Figure 1:**
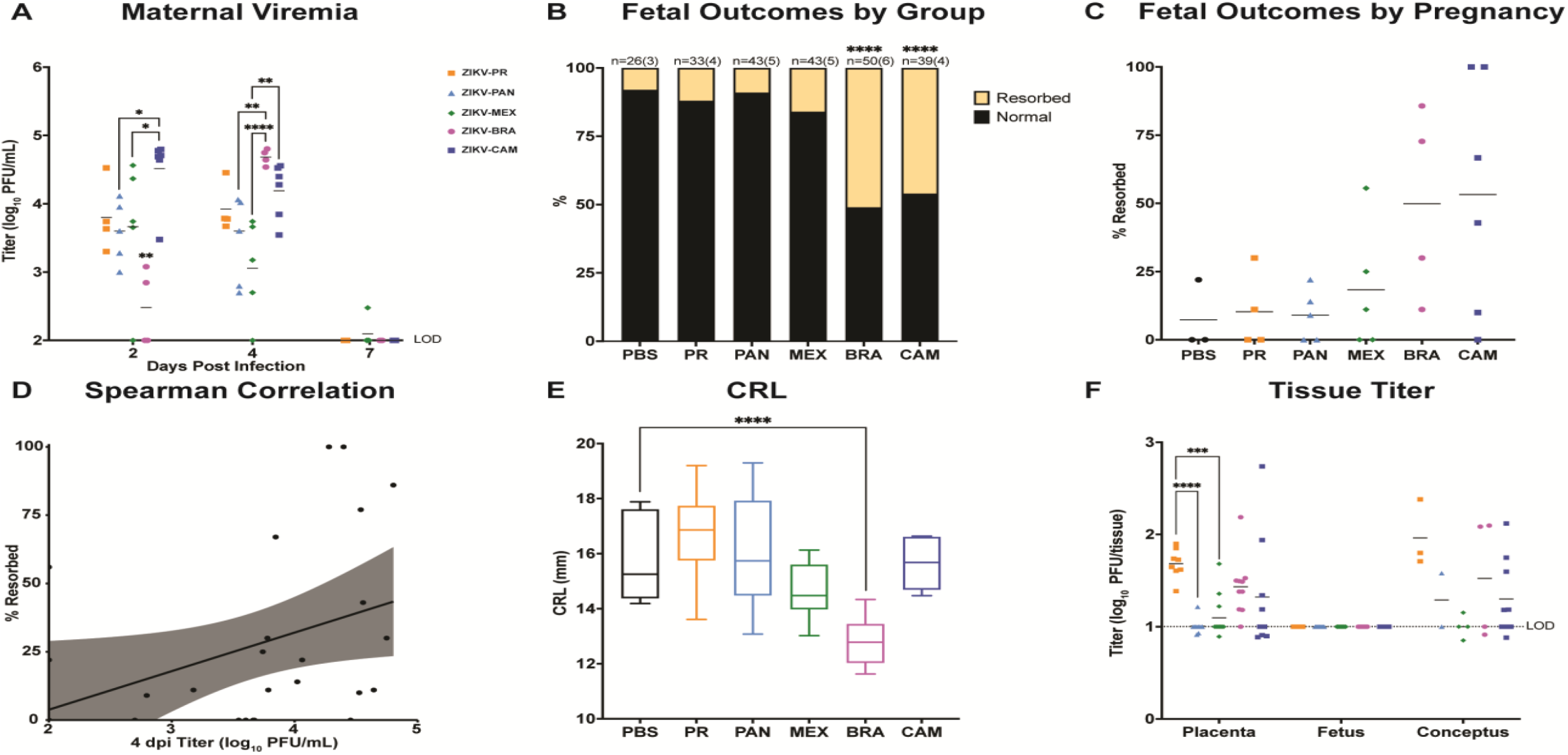
ZIKV strains are phenotypically heterogeneous. **(A)** Time-mated *Ifnar1*^−/−^ dams were inoculated with 10^3^ PFU of ZIKV on E7.5 and maternal infection was assessed by plaque assay on 2, 4, and 7 days post inoculation, and significance was determined by one-way ANOVA. **(B)** Rate of normal (black) vs. resorbed (yellow) fetuses at E14.5 after maternal infection at E7.5. Data presented are n = number of individual fetuses from (3-6 litters per treatment group). Significance was determined by Fisher’s exact test. **(C)** Pregnancy outcomes of individual animals in each treatment group. Data are presented as percent of fetuses resorbed in each pregnancy. **(D)** Spearman correlation of maternal serum titer at 4 dpi vs. resorption rate (r = 0.5200, p-value = 0.0054). **(E)** Crown-to-rump length (CRL) measurements in mm of morphologically normal fetuses at E14.5 using ImageJ software. Significance was determined by one-way ANOVA. **(F)** Tissue titer was measured by plaque assay for individual homogenized placentas, fetuses, and concepti (when the fetus and placenta were indistinguishable due to severe resorption). Symbols represent individual placenta, fetus, or conceptus from 4–6 independent experiments for each treatment group. Bars represent the mean viral titer of each treatment group and significance was determined by one-way ANOVA. Significance annotations for all figures: ****p≤0.0001; ***p≤0.001; **p≤0.01; *p≤0.05.

### Adverse fetal outcomes are dependent on the infecting ZIKV strain

Next, to assess fetal outcomes, dams were necropsied on E14.5. Gross examination of each conceptus revealed overt differences among fetuses within pregnancies and with uninfected counterparts. Fetuses appeared as either morphologically normal or undergoing embryo resorption, as defined in (20). At time of necropsy, we observed high rates of resorption from ZIKV-BRA- and ZIKV-CAM-infected pregnancies (ZIKV-BRA: 51% and ZIKV-CAM: 46%), which were significantly higher than the other virus-inoculated groups and PBS-inoculated controls (Fisher’s Exact test, *p* < 0.0001) (**Figure 1B**). In contrast, the proportion of resorbed fetuses for ZIKV-PR-, ZIKV-PAN-, and ZIKV-MEX-infected pregnancies did not differ significantly from each other or from PBS-inoculated controls (ZIKV-PR: 12%, ZIKV-PAN: 9%, and ZIKV-MEX: 16%, PBS: 8%, *p* > 0.1264) (**Figure 1B**). The rate of embryo resorption also varied between individual pregnancies within each treatment group (**Figure 1C**). Maternal viremia at 4 dpi positively correlated with increased fetal resorption across all virus groups (Spearman, *p* = 0.0111) (**Figure 1D**), but this trend was not observed within individual virus groups (Spearman, *p* > 0.1333). Therefore, our results demonstrate that multiple ZIKV genotypes differ in their propensity to cause fetal harm in this experimental model, and additional factors, beyond maternal infection, may contribute to fetal outcome.

### Fetal growth restriction is only evident following ZIKV-BRA infection

To further characterize pathogenic outcomes during pregnancy, we measured crown-to-rump length (CRL) to assess overall fetal growth (20, 22). Only fetuses that appeared morphologically normal were included for measurement of CRL to provide evidence for intrauterine growth restriction (IUGR). There was a statistically significant reduction in CRL in ZIKV-BRA fetuses compared to fetuses from PBS-inoculated dams (One-way ANOVA with Tukey’s multiple comparisons, *p* < 0.0001) (**Figure 1E**). In contrast, mean CRL did not differ significantly between fetuses from ZIKV-MEX-, ZIKV-CAM-, ZIKV- PAN-, ZIKV-PR-, and PBS-inoculated dams (*p* > 0.1561). The lack of apparent IUGR for the other ZIKV strains is contrary to other studies using Asian-lineage ZIKVs (French Polynesia and Cambodia) in which fetuses developed severe IUGR (22, 23). However, the discrepancy in outcomes may be the result of differences in timing of challenge and necropsy, dose and/or route of inoculation, dam age, litter size, or metrics for defining grossly normal fetuses compared to those undergoing resorption at a later embryonic age.

### No evidence for vertical transmission in any virus treatment groups

Next, to determine the potential of each ZIKV strain to be vertically transmitted, a subset of placentas and fetuses were collected for plaque assay at time of necropsy from each litter in all treatment groups. No infectious virus was detected by plaque assay in any fetus sample from any treatment group (**Figure 1F**) and the absence of ZIKV fetal infection was confirmed by RNA In Situ Hybridization (ISH). In contrast, virus was detected in placentas from all virus-inoculated groups at time of necropsy at E14.5 (7 dpi). ZIKV-PR placenta titers were significantly higher than ZIKV-PAN and ZIKV-MEX titers (One-way ANOVA with Tukey’s multiple comparisons, *p* <0.0001 and *p* = 0.0006), but only modestly higher than ZIKV-CAM and ZIKV-BRA titers (*p* = 0.5591 and *p* = 0.5693) (Figure 1F). In ZIKV-PR, ZIKV-PAN, ZIKV-MEX, and ZIKV-BRA groups, placenta titer was not a predictor of partner fetus outcome (One-way ANOVA, *p* > 0.0970). Although limited by the number of data points, ZIKV-CAM placentas with resorbed fetuses had significantly higher titers than those from normal fetuses (*p* = 0.0182). Therefore, we hypothesized that additional factors, outside of fetus and placenta virus levels, contribute to poor fetal outcomes.

### Severe placenta histopathological changes were consistently detected in ZIKV-CAM infected mice

To better characterize the impact of in-utero infection of different ZIKV strains, placental tissues were examined microscopically. In PBS-inoculated dams, we observed normal decidua, junctional zone, and labyrinth with normal maternal and fetal blood spaces (**Figure 2**). In contrast, ZIKV-inoculated dams displayed varying degrees of placental pathology, similar to what we have reported previously, with the most severe effects predominantly observed in the labyrinth zone, including necrosis, calcifications, thrombi, inflammation, and apoptosis (20, 21). Interestingly, the overall severity observed within virus groups was relatively subtle compared to our previous studies (20, 21), which may account for the higher background scores noted in the PBS control group (**Figure 2A-C**). There also were clear strain-specific differences in the amount of placental pathology observed, with ZIKV-CAM displaying the most severe histologic phenotype (**Figure 2**). Similar to placenta titer, pathology severity score was not a predictor of adverse fetal outcome for any treatment group.

**Figure 2:**
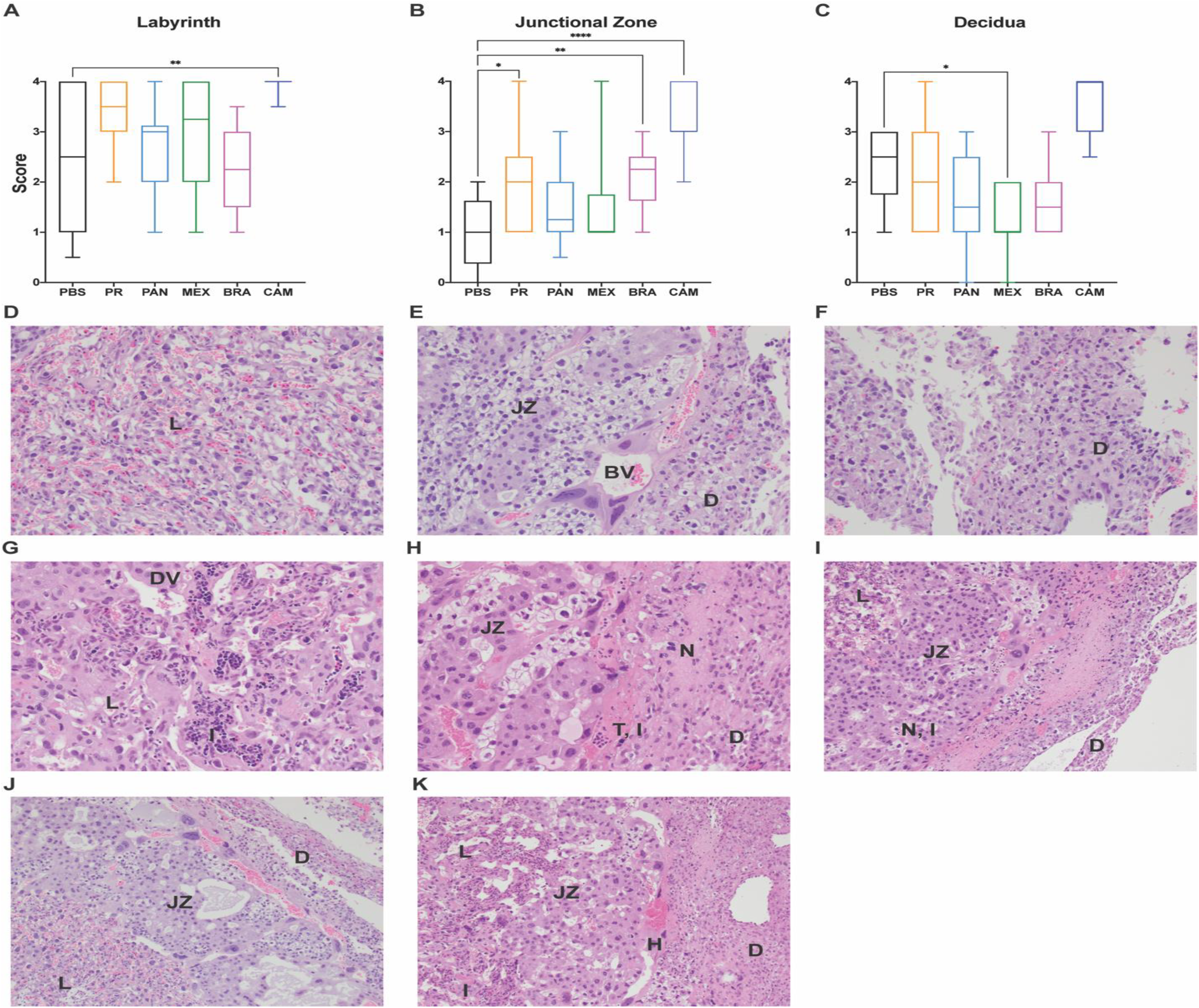
Placenta histopathology is dependent on infecting virus strain. **(A-C)** The degree of placenta pathology was rated on a scale of 0-4: zero represents normal histologic features and 4 represents the most severe features observed. Each zone of the placenta was scored individually for general overall pathology, amount of inflammation, and amount of vascular injury. Only ‘General’ scores are shown because they are representative of ‘inflammation’ and ‘vascular injury’ categories and do not differ significantly from ‘general’. Box and whiskers represent the minimum-to-maximum of all data points around the median. Data are representative of 3-6 independent animals for each treatment group. ****p≤0.0001; **p≤0.01; *p≤0.05 (Kruskal-Wallis ANOVA). **(D-F)** Normal histologic features of each placental zone - labyrinth (L), junctional zone and decidua (JZ), and decidua (D) - from PBS-inoculated dams. **(E)** Normal blood vessel (BV) between the junctional zone and decidua. **(G-I)** Severe histopathologic injury patterns from placentas from ZIKV-CAM-inoculated dams. **(G)** Dilated vasculature (DV) and inflammation (I) in the labyrinth. **(H)** Thrombus and inflammation (T & I) at the junction of the junctional zone and decidua, and necrosis (N) in the decidua. **(I)** Necrosis and inflammation (N & I) at the junction of the junctional zone and decidua, and necrosis of the decidua. **(J)** Normal pathology of labyrinth, junctional zone, and decidua from a PBS placenta **(K)** Severe pathology of labyrinth, junctional zone, and decidua from a ZIKV-CAM placenta. Dilated vascular space and inflammation in the labyrinth, hemorrhage (H) in the junctional zone, and necrosis in the decidua.

### ISG transcript abundance is elevated in placentas after ZIKV infection

Due to the lack of vertical transmission and an association between fetal outcome and placenta infection and pathology, we hypothesized that IFN induction in the placenta was responsible for determining fetal outcome. Indeed, it has previously been shown that type I IFN signaling, not the levels of virus, mediated pathology following intravaginal ZIKV infection in *Ifnar1*^+/−^ fetuses and placentas (23). Accordingly, we examined the transcriptional changes of the interferon-stimulated genes (ISGs) *Oasl2*, *Mx1*, and *Ifit1* in the placenta to determine if IFN induction (or the lack thereof) may be contributing to fetal demise. We observed that *Oasl2*, *Mx1*, and *Ifit1* were induced regardless of infecting ZIKV genotype in our model (**Figure 3A-C**). Interestingly, ZIKV-MEX placentas had significantly lower *Mx1*, *Oasl2*, and *Ifit1* transcript abundance compared to the other virus groups (One-way ANOVA, *p* < 0.0357). Across virus groups, pregnancies with better outcomes (i.e., lower rates of resorption) had higher expression of *Mx1* (Spearman, *p* = 0.0169) (Figure 1D), and ZIKV-CAM placentas with normal fetuses expressed higher levels of *Mx1* than their resorbed counterparts (*p* = 0.0464; mean ± SEM: −18.01 ± 7.650; n = 10). However, there was no correlation between ISG expression and pregnancy outcome within any virus group (Spearman, *p* > 0.0833). Also, ISG expression and placenta histopathology scores showed no clear relationship. Across virus groups, maternal serum titer at 4 dpi positively correlated with increased expression of *Oasl2*, *Mx1*, and *Ifit1* in the placenta (Spearman, *p* < 0.0487) (**Figure E-G**). These data suggest a neutral, or modestly protective, role for the IFN response in our model.

**Figure 3:**
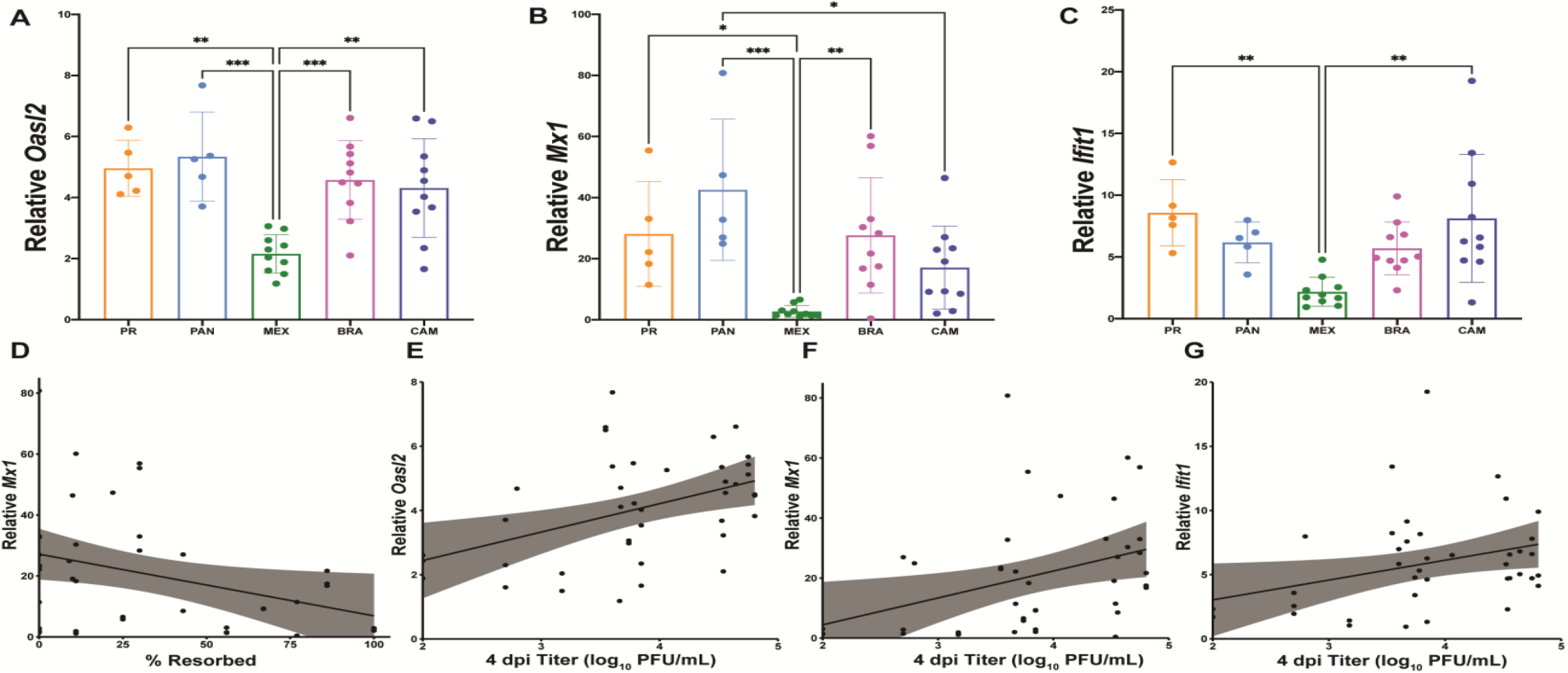
ISG expression is elevated in placentas from ZIKV-infected dams. RNA was extracted from placentas harvested on E14.5 and expression of the ISGs *Oasl2* **(A)**, *Mx1* **(B)**, *Ifit1* **(C)** were analyzed by QPCR. Levels were normalized to *Gapdh* and then ddCT was calculated relative to placentas harvested from PBS-inoculated dams. 1-4 placentas from 2 litters of PBS-inoculated and 4-5 litters of each ZIKV treatment group were analyzed. Mean with standard deviation are shown. ***p≤0.001; **p≤0.01; *p≤0.05 (one-way ANOVA). **(D-G)** Spearman correlations with shaded 95% confidence interval are shown for % resorbed vs. relative *Mx1* expression (r = −0.6593; p-value = 0.0169) **(D)**, and 4 dpi maternal titer vs. relative expression of *Oasl2* (r = 0.3752; p-value = 0.0171) **(E)**, *Mx1* (r = 0.4111; p-value = 0.0084) **(F)**, and *Ifit1* (r = 0.3137; p-value = 0.0487) **(G)**.

We then examined if infection outcomes were due to differential expression of IFN-λ or its heterodimeric receptor *Ifnλr1/Il10rβ*. We measured the relative transcript abundance of *Ifnλr1*, *Ifnλ2*, and *Ifnλ3* in the placenta at time of necropsy at E14.5. *Ifnλ1* is a pseudogene and the genomic region of *Ifnλ4* is missing in mice (28) and, therefore, were not measured. Consistent with our ISG data, we observed modest induction of *Ifnλr1* for all ZIKV strains (**Figure 4A**). Importantly, pregnancies with lower rates of resorption had higher expression of *Ifnλr1* (Spearman, *p* = 0.0302) (**Figure 4B**), and ZIKV-CAM placentas with normal fetuses had higher expression of *Ifnλr1* than their resorbed counterparts (*p* = 0.0087; mean ± SEM: −1.148 ± 0.333; n = 10). In contrast, *Ifnλ2* and *Ifnλ3* were not induced in any placenta sample from ZIKV-infected mice (**Figure 4C-D**). This, perhaps, was not surprising since a previous mouse study showed that type III IFNs played little to no role in placental antiviral defenses before placentation (25). In our model, dams were infected on E7.5 and placental development is not complete until E8.5-10.5 in mice (29, 30). Still, it remains unknown whether the mouse placenta constitutively releases type III IFNs in a manner similar to the human placenta, or whether these IFNs are induced systemically or in response to placental infection (31). Here, we did not detect robust evidence for induction of type III IFNs despite detection of infectious virus in the placenta at time of necropsy at E14.5.

**Figure 4:**
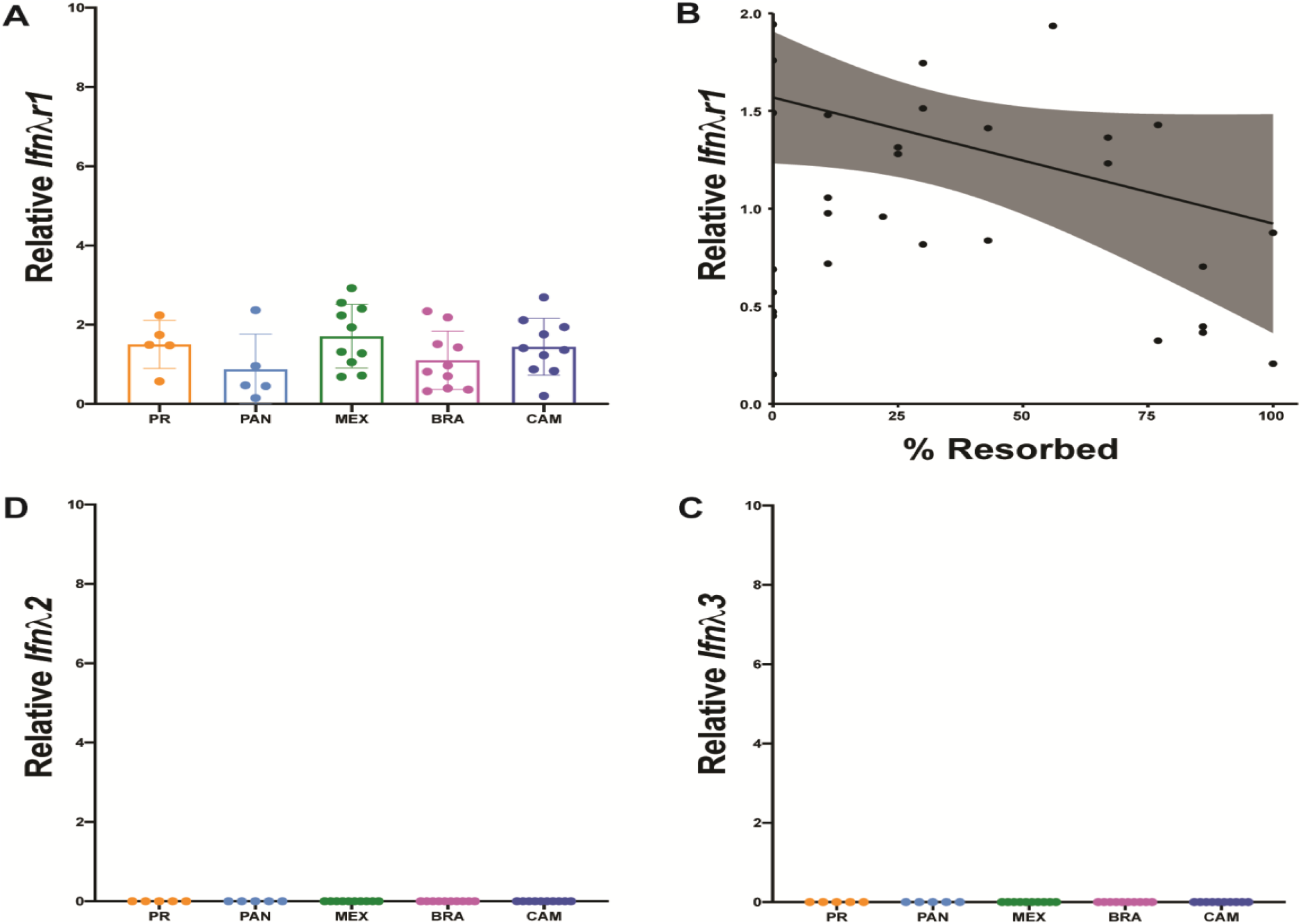
No robust evidence for Type III IFN induction. RNA was extracted from placentas harvested on E14.5 and relative expression was analyzed by QPCR. Levels were normalized to *Gapdh* and then ddCT was calculated relative to placentas harvested from PBS-inoculated dams. **(A)** *Ifnλr1* expression. **(B)** Spearman correlation of % resorbed and *Ifnλr1* expression from individual pregnancies (r = −0.6099; p-value = 0.0302). Relative expression of IFNλ2 **(C)** and IFNλ3 **(D)**.

## DISCUSSION

By comparing five ZIKV strains representing the viral genetic diversity in the Americas (32), we provide experimental evidence that there is strain-dependent phenotypic heterogeneity in pregnancy outcomes following in-utero ZIKV exposure. In our pregnant *Ifnar1^−/−^* mouse model, ZIKV-CAM and ZIKV-BRA caused significantly more embryo resorption. Maternal infection with ZIKV-PAN and ZIKV-MEX resulted in low levels of placenta infection with varying degrees of placenta pathology, and overall low rates of embryo resorption. In contrast, ZIKV-PR replicated to high titers in the placenta that corresponded to severe histopathology but did not result in fetal demise. No strain resulted in detectable fetal infection, which is different from what we have reported previously with African-lineage ZIKV (20, 21). However, the absence of ZIKV fetal infection does not preclude the possibility that pathology may develop later in pregnancy or even postnatally, similar to what has been observed in humans (33). It is unknown whether adverse pregnancy outcomes require direct infection of the fetus (i.e., vertical transmission) or whether pathophysiology at the maternal-fetal interface (MFI) without vertical transmission is sufficient to cause adverse outcomes. Placental insufficiency is now being recognized as a potential contributor to some of these adverse outcomes (34, 35), and our data suggest that pregnancy loss is not solely driven by fetal infection.

One possible explanation for differences in fetal outcomes observed between treatment groups could be due to differences in activation of and/or susceptibility to antiviral signaling at the MFI. It is becoming increasingly apparent that IFN responses can have protective and/or pathogenic effects in pregnancy (reviewed in (28)). Protection associated with IFN production prevents uncontrolled virus replication, fetal infection, and maternal mortality (36–38); however, overproduction of type I IFNs are known to be an underlying cause of pregnancy complications, including developmental defects similar to those that result from infections with teratogenic pathogens (23, 39, 40). As a result, there likely is a critical balance that must occur between the beneficial antiviral effects of the IFN response to virus infections during pregnancy and the pathological consequences that may result from excessive production of IFNs. We examined the relative levels of the ISGs *Oasl2*, *Mx1*, and *Ifit1* in the placenta because of their known relevance to mouse (23, 41–44) and nonhuman primate (45) models of ZIKV infection, broad-spectrum antiviral functions (46–50), contributions to placental pathology (39, 51), and general involvement in the success of human pregnancies (28, 36, 52). Our data suggest that IFN activation did not contribute to fetal demise and, in some cases, may have played a protective role. Maternal viremia appeared to drive ISG induction in the placenta, which may not be surprising given that the mouse labyrinth is perfused with maternal blood (53). Higher maternal viremia also positively correlated with increased resorption rate across virus groups. Therefore, maternal viremia may also contribute to an increased risk of adverse fetal outcomes, alone or in combination with IFN-dependent causes, direct pathogenic effects of the virus, or as a bystander effect associated with immune responses unrelated to IFN induction. Importantly, we only assessed ISG expression at a single timepoint, at time of necropsy (E14.5, 7 dpi), and expression profiles may differ depending on the timing of collection. More studies are needed to better understand antiviral signaling at the MFI and the mechanisms these virus strains exploit to harm the feto-placental unit.

Differences observed in fetal outcomes and histopathology across ZIKV strains may also be due to virus genetic determinants of virulence and pathogenesis during congenital infection. Because contemporary ZIKV isolates are so closely related, they are oftentimes used interchangeably in laboratory research. But even though there is high genetic similarity between ZIKV strains, it is possible that subtle genotypic differences could result in small, but biologically important, phenotypic differences between strains. For example, evidence suggests that ZIKV virulence can be governed both by viral nucleotide sequence and/or amino acid sequence (41, 54–58), but the impact of a single amino acid substitution may vary in the different strains chosen for analyses (20, 59). ZIKV strains used here share >98% genome-wide nucleotide identity and while it is unclear whether the differences in amino acid and/or nucleotide sequence are responsible for the differences in the observed phenotypes, it is possible that there is not a single determinant of ZIKV fetal pathogenicity. Future reverse genetic studies will be needed to fully understand if there is a link between viral genotype and phenotype.

Our findings highlight that phenotypic heterogeneity exists between closely related ZIKV strains that are commonly used for pathogenesis studies. To more rigorously assess the relative capacity of Asian/American-lineage ZIKVs to cause adverse fetal outcome, future studies should carefully consider the specific characteristics of the virus strains being used and consider them in the specific context of the questions being asked. One important limitation to our study is that it is unclear whether the same phenotypes would be recapitulated during human infection. Further, we do not argue that the phenotypic differences we observe between strains indicate diminished risk of adverse outcomes following infection during pregnancy with a certain ZIKV genotype (60). On the contrary, the presence of infectious ZIKV in the placenta for all strains tested is concerning and suggests that all ZIKV strains have the capacity to harm the developing fetus depending on the specific pathophysiological context of infection at the MFI. Here, our results provide a comparative framework to further investigate underlying factors that determine fetal outcome during in-utero ZIKV exposure.

## MATERIALS & METHODS

### Ethical Approval

This study was approved by the University of Minnesota, Twin-Cities Institutional Animal Care and Use Committees (Animal Care and Use Protocol Number 1804-35828).

### Cells and Viruses

African Green Monkey kidney cells (Vero; ATCC #CCL-81) were maintained in Dulbecco’s modified Eagle medium (DMEM) supplemented with 10% fetal bovine serum (FBS; Corning, Manassas, VA), 1X Antibiotic Antimycotic solution (Corning, Manassas, VA) and incubated at 37°C in 5% CO_2_. *Aedes albopictus* mosquito cells (C6/36; ATCC #CRL-1660) were maintained in DMEM supplemented with 10% fetal bovine serum (FBS; Hyclone, Logan, UT), 2 mM L-glutamine, 1.5 g/L sodium bicarbonate, 1X Antibiotic Antimycotic solution, and incubated at 28°C in 5% CO_2_. The cell lines were obtained from the American Type Culture Collection, were not further authenticated, and were not specifically tested for mycoplasma.

ZIKV strain PRVABC59 (ZIKV-PR; GenBank:KU501215) was originally isolated from a traveler to Puerto Rico in 2015 with three rounds of amplification on Vero cells. ZIKV strain R116265 (ZIKV-MEX; GenBank:KX766029) was originally isolated from a 73-year-old-male traveling in Mexico in 2016 with a single round of amplification on Vero cells (CDC, Ft. Collins, CO). ZIKV strain 259249 (ZIKV-PAN; GenBank:KX156775) was originally isolated from a human serum sample from Panama in 2015 with two rounds of amplification on Vero cells, followed by one round of amplification on C6/36 mosquito cells. ZIKV strain FSS13025 (ZIKV-CAM; GenBank:JN860885) was originally isolated from a child in Cambodia with three rounds of amplification on Vero cells. Master stocks were obtained from Brandy Russell (CDC, Ft. Collins, CO). ZIKV strain Paraiba_01 (ZIKV-BRA; GenBank:KX280026) was originally isolated from human serum in Brazil in 2015 with two rounds of amplification on Vero cells, and a master stock was obtained from Dr. Kevin Noguchi at Washington University in St. Louis (St. Louis, MO). Virus challenge stocks were prepared by inoculation onto a confluent monolayer of C6/36 mosquito cells. We deep sequenced our virus stocks to verify the expected origin (see next section for details).

### Deep Sequencing

A vial of all viral stocks used for challenges were each deep sequenced by preparing libraries of fragmented double-stranded cDNA using methods similar to those previously described (20, 21, 61). Briefly, the sample was centrifuged at 5000 rcf for five minutes. The supernatant was then filtered through a 0.45-μm filter. Viral RNA was isolated using the QIAamp MinElute Virus Spin Kit (Qiagen, Germantown, MD), omitting carrier RNA. Eluted vRNA was then treated with DNAse I. Double-stranded DNA was prepared with the Superscript Double-Stranded cDNA Synthesis kit (Invitrogen, Carlsbad, CA) and priming with random hexamers. Agencourt Ampure XP beads (Beckman Coulter, Indianapolis, IN) were used to purify double-stranded DNA. The purified DNA was fragmented with the Nextera XT kit (Illumina, Madison, WI), tagged with Illumina-compatible primers, and then purified with Agencourt Ampure XP beads. Purified libraries were then sequenced with 2 × 300 bp kits on an Illumina MiSeq.

### Sequence Analysis

Viral stock sequences were analyzed using a modified version of the viral-ngs workflow developed by the Broad Institute (http://viral-ngs.readthedocs.io/en/latest/description.html) implemented in DNANexus and using bbmap local alignment in Geneious Pro (Biomatters, Ltd., Auckland, New Zealand). Briefly, using the viral-ngs workflow, host-derived reads that map to a human sequence database and putative PCR duplicates were removed. The remaining reads were loaded into Geneious Pro and mapped to NCBI Genbank Zika (GenBank:KX601166) reference sequences using bbmap local alignment. Mapped reads were aligned using Geneious global alignment and the consensus sequence was used for intra sample variant calling. Variants were called that fit the following conditions: have a minimum *p*-value of 10e-60, a minimum strand bias of 10e-5 when exceeding 65% bias, and were nonsynonymous. Consensus-level nucleotide substitutions and minor nucleotide variants are reported in **Table 3.**

**Table 3:**
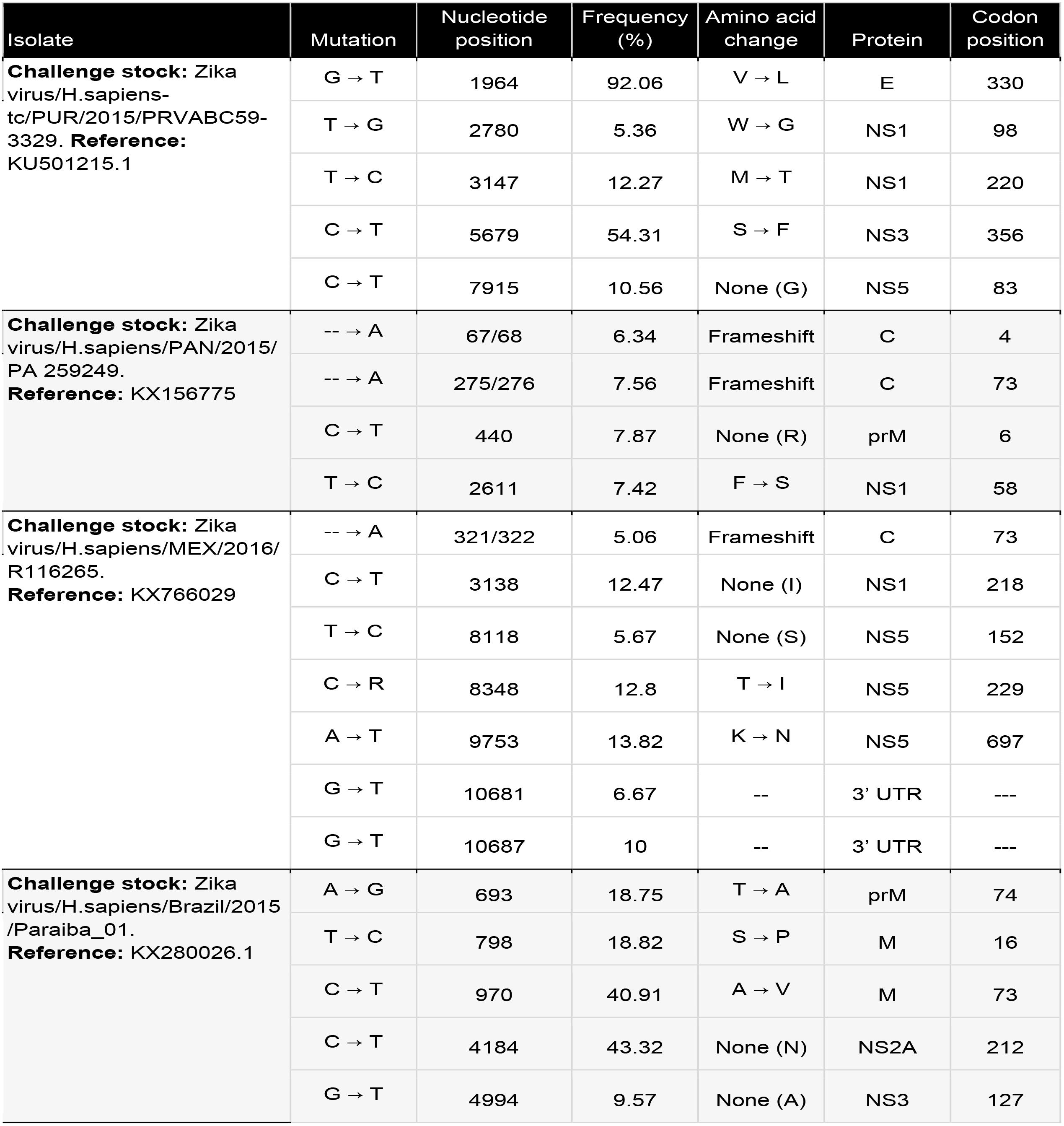
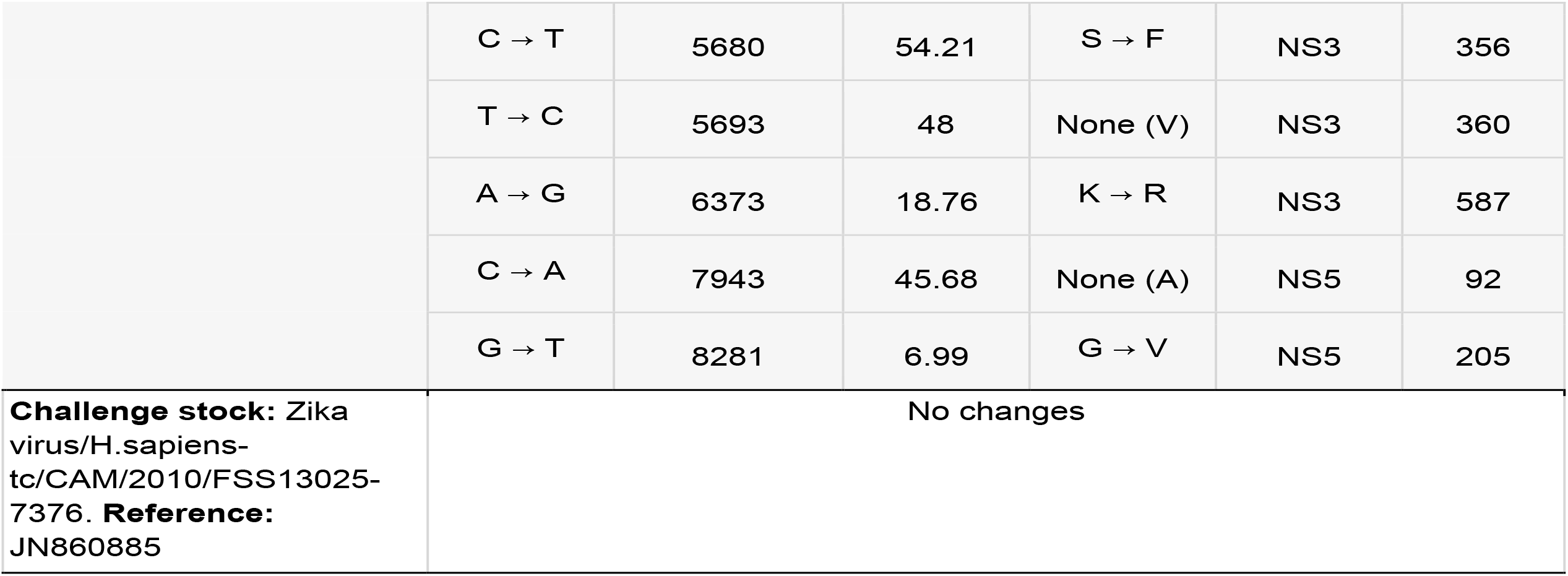
Nucleotide variants in challenge stocks relative to the GenBank reference sequence. Only variants found in >50% of sequences are shown.

### Plaque assay

Quantification of virus titer in maternal serum, placenta, and fetuses were completed by plaque assay on Vero cells. Duplicate wells were infected with 0.1 mL aliquots from serial 10-fold dilutions in growth medium and virus was adsorbed for 1 hour. After incubation, the monolayers were overlaid with 3 mL containing a 1:1 mixture of 1.2% oxoid agar and 2X DMEM (Gibco, Carlsbad, CA) with 10% (vol/vol) FBS and 2% (vol/vol) Antibiotic Antimycotic. Cells were incubated at 37°C in 5% CO_2_ for three days (ZIKV-PR, ZIKV-BRA, ZIKV-CAM), four days (ZIKV-PAN), or five days (ZIKV-MEX) for plaque development. Cell monolayers were then stained with 3 mL of overlay containing a 1:1 mixture of 1.2% oxoid agar with 4% neutral red (Gibco) and 2X DMEM with 2% (vol/vol) FBS, and 2% (vol/vol) Antibiotic Antimycotic. Cells were incubated overnight at 37°C in 5% CO_2_ and plaques were counted.

### Mice

Female *Ifnar1*^−/−^ mice on the C57BL/6 background were bred in the specific pathogen-free animal facilities of the University of Minnesota within the College of Veterinary Medicine. Male C57BL/6 were purchased from Jackson Laboratories. Timed matings between female *Ifnar1^−/−^* mice and male C57BL/6 mice resulted in *Ifnar1*^+/−^ progeny.

### Subcutaneous inoculation

All pregnant dams were between six and ten weeks of age and were randomly assigned to infected or control groups. Matings between *Ifnar1^−/−^* dams and wildtype sires were timed by checking for the presence of a vaginal plug, indicating gestational age E0.5. At embryonic day 7.5 (E7.5) dams were inoculated in the right hind footpad with 1×10^3^ PFU of the selected ZIKV strain in sterile PBS or with sterile PBS alone to serve as experimental controls. All animals were closely monitored by laboratory staff for adverse reactions and/ or clinical signs of disease. A submandibular blood draw was performed at 2, 4, and 7 days post inoculation (dpi), and serum was collected to verify viremia. Mice were humanely euthanized and necropsied at E14.5.

### Mouse necropsy

Following inoculation with ZIKV or PBS, mice were sacrificed at E14.5. Tissues were carefully dissected using sterile instruments that were changed between each mouse to minimize possible cross contamination. Each organ and neonate were morphologically evaluated in situ prior to removal. Using sterile instruments, the uterus was removed and dissected to remove individual concepti. Each conceptus was placed in a sterile culture dish and dissected to separate the fetus and the placenta, when possible, for gross evaluation. Fetuses were characterized as “normal” or “resorbed”, with the latter being defined as having significant growth retardation and reduced physiological structure compared to littermates and controls, accompanied by clearly evident developmental delay or visualization of a macroscopic plaque in the uterus. A subset of fetuses and placentas from each litter were reserved for viral titer analysis (preserved in PBS supplemented with 20% FBS and 1% Antibiotic Antimycotic) or fixed in 10% neutral buffered formalin for imaging and histology.

### Crown-to-rump length

Crown-to-rump length (CRL) was measured by tracing the distance from the crown of the head to the base of the tail, using ImageJ. Infection-induced resorbed fetuses were excluded from measurement analyses because they would not survive if the pregnancy was allowed to progress to term (20).

### Histology

Placenta tissues were fixed in 10% neutral buffered formalin at room temperature for 36-48 hours and then transferred to 70% ethanol until alcohol-processed and embedded in paraffin. Paraffin sections (5 μm) were stained with hematoxylin and eosin (H&E) and the degree of pathology was scored by a blinded pathologist, as described in (20). The degree of placental pathology was rated on a relative scale of 0–4: zero represents normal histologic features and 4 represents the most severe features observed. Each zone of the placenta was scored individually for general overall pathology, amount of inflammation, and amount of vascular injury. Only ‘General’ scores are shown because they were representative of ‘inflammation’ and ‘vascular injury’ scores.

### In situ hybridization

Immediately following necropsy, fetuses were fixed in 10% neutral buffered formalin at room temperature for 36-48 hours and then transferred to 70% ethanol until alcohol-processed and embedded in paraffin. Paraffin sections (5 μm) were deparaffinized and a hydrogen peroxide quench was performed, followed by boiling in target retrieval reagent (catalog #322000). Tissue was then incubated in Protease Plus solution (catalog #322330) in a HybEZ II Oven at 40°C before hybridization with the ZIKV probe (catalog #468361) and chromogen labeling using the RNAscope 2.5 HD Red Assay (catalog #322360). In Situ Hybridization (ISH) was performed using the RNAscope Assay using products and instructions (62) provided by the manufacturer (Advanced Cell Diagnostics. Inc., Newark, CA). Each ISH run included ZIKV-infected positive control tissue to confirm the protocol was run as properly. After labeling, tissue was counterstained using hematoxylin before cover-slipping for evaluation.

### Fetal and Placental viral titers

An Omni TH115 Homogenizer (Omni International, Omni Tissue Homogenizer (TH) - 115V) was used to homogenize fetus and placenta samples following necropsy. Samples were submerged in chilled PBS supplemented with 20% FBS and 1% Antibiotic Antimycotic in 15mL Omni sealed plastic tubes (Omni International, Catalog # 00-2015-25). Omni soft tissue probes (Omni International, Catalog # 30750) were used to homogenize samples at the highest speed for 15 seconds (placentas) or 30 seconds (fetuses). Homogenized samples were clarified by centrifugation at 10,000 x g for 2 minutes. The supernatant was removed and 0.1mL was immediately plated for plaque assay. The remainder was stored at −80°C.

### Innate immune gene RT-QPCR in mouse placenta

RNA was extracted and purified from placentas using a Direct-zol RNA kit (Zymo Research). The High-Capacity RNA-to-cDNA Kit (Applied Biosystems) was used to synthesize cDNA. Quantitative polymerase chain reaction (qPCR) using PowerUp SYBR Green Master Mix (Applied Biosystems) was used to quantify innate immune genes and run on a QuantStudio 3 (Applied Biosystems). The following PrimeTime Primers (Integrated DNA Technologies) were used: *Ifnλr1:* Mm.PT.58.10781457, *Ifnλ2:* Mm.PT.58.31485549, *Ifnλ3:* Mm.PT.58.8956530, *Ifit1:* Mm.PT.58.32674307, *Mx1:* Mm.PT.58.12101853.g, *Oasl2:* Mm.PT.56a.17167264, and *Gapdh:* Mm.PT.39a.1. Innate immune genes were normalized to *Gapdh* and then 2-delta delta CT was calculated relative to PBS-inoculated controls.

### Data analysis

All analyses were performed using GraphPad Prism. Unpaired Student’s t-test was used to determine significant differences in crown-rump lengths. Fisher’s exact test was used to determine differences in rates of normal versus resorbed concepti. One-way ANOVA with Tukey’s multiple comparison test was conducted to compare virus titers in maternal serum, placentas, fetuses, and concepti. Nonparametric Spearman correlation was used to evaluate the relationship between variables.

## Data availability

Virus stock sequence data have been deposited in the Sequence Read Archive (SRA) with accession codes SRX4510825, SRR14467422, and SRR14467421. The authors declare that all other data supporting the findings of this study are available within the article.

## ACKNOWLEDGEMENTS

We thank the University of Minnesota, Twin Cities Comparative Pathology Shared Resource for preparation of histological sections. Research reported in this publication was supported by the National Institute of Allergy and Infectious Diseases of the National Institutes of Health under Award Number R01AI132563 to M.T.A. A.S.J is supported by the University of Minnesota, Twin Cities Institute for Molecular Virology Training Program predoctoral fellowship, T32AI083196. The publication’s contents are solely the responsibility of the authors and do not necessarily represent the official views of the NIH.

## REFERENCES

1. Ospina ML, Tong VT, Gonzalez M, Valencia D, Mercado M, Gilboa SM, Rodriguez AJ, Tinker SC, Rico A, Winfield CM, Pardo L, Thomas JD, Avila G, Villanueva JM, Gomez S, Jamieson DJ, Prieto F, Meaney-Delman D, Pacheco O, Honein MA. 2020. Zika Virus Disease and Pregnancy Outcomes in Colombia. N Engl J Med 383:537–545.

2. Sanchez Clemente N, Brickley EB, Paixão ES, De Almeida MF, Gazeta RE, Vedovello D, Rodrigues LC, Witkin SS, Passos SD. 2020. Zika virus infection in pregnancy and adverse fetal outcomes in São Paulo State, Brazil: a prospective cohort study. Sci Rep 10:12673.

3. Smoots AN, Olson SM, Cragan J, Delaney A, Roth NM, Godfred-Cato S, Jones AM, Nahabedian JF, Fornoff J, Sandidge T, Yazdy MM, Higgins C, Olney RS, Eckert V, Forkner A, Fox DJ, Stolz A, Crawford K, Cho SJ, Knapp M, Ahmed MF, Lake-Burger H, Elmore AL, Langlois P, Breidenbach R, Nance A, Denson L, Caton L, Forestieri N, Bergman K, Humphries BK, Leedom VO, Tran T, Johnston J, Valencia-Prado M, Pérez-González S, Romitti PA, Fall C, Bryan JM, Barton J, Arias W, St John K, Mann S, Kimura J, Orantes L, Martin B, de Wilde L, Ellis EM, Song Z, Akosa A, Goodroe C, Ellington SR, Tong VT, Gilboa SM, Moore CA, Honein MA. 2020. Population-Based Surveillance for Birth Defects Potentially Related to Zika Virus Infection - 22 States and Territories, January 2016-June 2017. MMWR Morb Mortal Wkly Rep 69:67–71.

4. Mulkey SB, Arroyave-Wessel M, Peyton C, Bulas DI, Fourzali Y, Jiang J, Russo S, McCarter R, Msall ME, du Plessis AJ, DeBiasi RL, Cure C. 2020. Neurodevelopmental Abnormalities in Children With In Utero Zika Virus Exposure Without Congenital Zika Syndrome. JAMA Pediatr 174:269–276.

5. Valdes V, Zorrilla CD, Gabard-Durnam L, Muler-Mendez N, Rahman ZI, Rivera D, Nelson CA. 2019. Cognitive Development of Infants Exposed to the Zika Virus in Puerto Rico. JAMA Netw Open 2:e1914061.

6. Stringer EM, Martinez E, Blette B, Toval Ruiz CE, Boivin M, Zepeda O, Stringer JSA, Morales M, Ortiz-Pujols S, Familiar I, Collins M, Chavarria M, Goldman B, Bowman N, de Silva A, Westreich D, Hudgens M, Becker-Dreps S, Bucardo F. 2021. Neurodevelopmental Outcomes of Children Following In Utero Exposure to Zika in Nicaragua. Clin Infect Dis 72:e146–e153.

7. Pimentel R, Khosla S, Rondon J, Peña F, Sullivan G, Perez M, Mehta SD, Brito MO. 2021. Birth Defects and Long-Term Neurodevelopmental Abnormalities in Infants Born During the Zika Virus Epidemic in the Dominican Republic. Ann Glob Health 87:4.

8. Aguilar Ticona JP, Nery N, Ladines-Lim JB, Gambrah C, Sacramento G, de Paula Freitas B, Bouzon J, Oliveira-Filho J, Borja A, Adhikarla H, Montoya M, Chin A, Wunder EA, Ballalai V, Vieira C, Belfort R, P Almeida AR, Reis MG, Harris E, Ko AI, Costa F. 2021. Developmental outcomes in children exposed to Zika virus in utero from a Brazilian urban slum cohort study. PLoS Negl Trop Dis 15:e0009162.

9. Musso D, Ko AI, Baud D. 2019. Zika Virus Infection - After the Pandemic. N Engl J Med 381:1444–1457.

10. Cauchemez S, Besnard M, Bompard P, Dub T, Guillemette-Artur P, Eyrolle-Guignot D, Salje H, Van Kerkhove MD, Abadie V, Garel C, Fontanet A, Mallet HP. 2016. Association between Zika virus and microcephaly in French Polynesia, 2013-15: a retrospective study. Lancet 387:2125–2132.

11. Rice ME, Galang RR, Roth NM, Ellington SR, Moore CA, Valencia-Prado M, Ellis EM, Tufa AJ, Taulung LA, Alfred JM, Pérez-Padilla J, Delgado-López CA, Zaki SR, Reagan-Steiner S, Bhatnagar J, Nahabedian JF, Reynolds MR, Yeargin-Allsopp M, Viens LJ, Olson SM, Jones AM, Baez-Santiago MA, Oppong-Twene P, VanMaldeghem K, Simon EL, Moore JT, Polen KD, Hillman B, Ropeti R, Nieves-Ferrer L, Marcano-Huertas M, Masao CA, Anzures EJ, Hansen RL, Pérez-Gonzalez SI, Espinet-Crespo CP, Luciano-Román M, Shapiro-Mendoza CK, Gilboa SM, Honein MA. 2018. Vital Signs: Zika-Associated Birth Defects and Neurodevelopmental Abnormalities Possibly Associated with Congenital Zika Virus Infection - U.S. Territories and Freely Associated States, 2018. MMWR Morb Mortal Wkly Rep 67:858–867.

12. Nogueira ML, Nery Júnior NRR, Estofolete CF, Bernardes Terzian AC, Guimarães GF, Zini N, Alves da Silva R, Dutra Silva GC, Junqueira Franco LC, Rahal P, Bittar C, Carneiro B, Vasconcelos PFC, Freitas Henriques D, Barbosa DMU, Lopes Rombola P, de Grande L, Negri Reis AF, Palomares SA, Wakai Catelan M, Cruz LEAA, Necchi SH, Mendonça RCV, Penha Dos Santos IN, Alavarse Caron SB, Costa F, Bozza FA, Soares de Souza A, Brandão de Mattos CC, de Mattos LC, Vasilakis N, Oliani AH, Vaz Oliani DCM, Ko AI. 2018. Adverse birth outcomes associated with Zika virus exposure during pregnancy in São José do Rio Preto, Brazil. Clin Microbiol Infect 24:646–652.

13. Ximenes RAA, Miranda-Filho DB, Montarroyos UR, Martelli CMT, Araújo TVB, Brickley E, Albuquerque MFPM, Souza WV, Ventura LO, Ventura CV, Gois AL, Leal MC, Oliveira DMDS, Eickmann SH, Carvalho MDCG, Silva PFSD, Rocha MAW, Ramos RCF, Brandão-Filho SP, Cordeiro MT, Bezerra LCA, Dimech G, Valongueiro S, Pires P, Castanha PMDS, Dhalia R, Marques-Júnior ETA, Rodrigues LC, Microcephaly Epidemic Research Group MERG. 2021. Zika-related adverse outcomes in a cohort of pregnant women with rash in Pernambuco, Brazil. PLoS Negl Trop Dis 15:e0009216.

14. Brasil P, Pereira JP, Moreira ME, Ribeiro Nogueira RM, Damasceno L, Wakimoto M, Rabello RS, Valderramos SG, Halai UA, Salles TS, Zin AA, Horovitz D, Daltro P, Boechat M, Raja Gabaglia C, Carvalho de Sequeira P, Pilotto JH, Medialdea-Carrera R, Cotrim da Cunha D, Abreu de Carvalho LM, Pone M, Machado Siqueira A, Calvet GA, Rodrigues Baião AE, Neves ES, Nassar de Carvalho PR, Hasue RH, Marschik PB, Einspieler C, Janzen C, Cherry JD, Bispo de Filippis AM, Nielsen-Saines K. 2016. Zika Virus Infection in Pregnant Women in Rio de Janeiro. N Engl J Med 375:2321–2334.

15. Barbeito-Andrés J, Schuler-Faccini L, Garcez PP. 2018. Why is congenital Zika syndrome asymmetrically distributed among human populations. PLoS Biol 16:e2006592.

16. Victora CG, Schuler-Faccini L, Matijasevich A, Ribeiro E, Pessoa A, Barros FC. 2016. Microcephaly in Brazil: how to interpret reported numbers. Lancet 387:621–624.

17. Miner JJ. 2017. Congenital Zika virus infection: More than just microcephaly. Sci Transl Med 9:eaan8195.

18. Adebanjo T, Godfred-Cato S, Viens L, Fischer M, Staples JE, Kuhnert-Tallman W, Walke H, Oduyebo T, Polen K, Peacock G, Meaney-Delman D, Honein MA, Rasmussen SA, Moore CA, Contributors. 2017. Update: Interim Guidance for the Diagnosis, Evaluation, and Management of Infants with Possible Congenital Zika Virus Infection - United States, October 2017. MMWR Morb Mortal Wkly Rep 66:1089–1099.

19. Lambrechts L. 2021. Did Zika virus attenuation or increased virulence lead to the emergence of congenital Zika syndrome. J Travel Med taab041.

20. Jaeger AS, Murrieta RA, Goren LR, Crooks CM, Moriarty RV, Weiler AM, Rybarczyk S, Semler MR, Huffman C, Mejia A, Simmons HA, Fritsch M, Osorio JE, Eickhoff JC, O’Connor SL, Ebel GD, Friedrich TC, Aliota MT. 2019. Zika viruses of African and Asian lineages cause fetal harm in a mouse model of vertical transmission. PLoS Negl Trop Dis 13:e0007343.

21. Jaeger AS, Weiler AM, Moriarty RV, Rybarczyk S, O’Connor SL, O’Connor DH, Seelig DM, Fritsch MK, Friedrich TC, Aliota MT. 2020. Spondweni virus causes fetal harm in Ifnar1^−/−^ mice and is transmitted by Aedes aegypti mosquitoes. Virology 547:35–46.

22. Miner JJ, Cao B, Govero J, Smith AM, Fernandez E, Cabrera OH, Garber C, Noll M, Klein RS, Noguchi KK, Mysorekar IU, Diamond MS. 2016. Zika Virus Infection during Pregnancy in Mice Causes Placental Damage and Fetal Demise. Cell 165:1081–1091.

23. Yockey LJ, Jurado KA, Arora N, Millet A, Rakib T, Milano KM, Hastings AK, Fikrig E, Kong Y, Horvath TL, Weatherbee S, Kliman HJ, Coyne CB, Iwasaki A. 2018. Type I interferons instigate fetal demise after Zika virus infection. Sci Immunol 3:eaao1680.

24. Aubry F, Jacobs S, Darmuzey M, Lequime S, Delang L, Fontaine A, Jupatanakul N, Miot EF, Dabo S, Manet C, Montagutelli X, Baidaliuk A, Gámbaro F, Simon-Lorière E, Gilsoul M, Romero-Vivas CM, Cao-Lormeau VM, Jarman RG, Diagne CT, Faye O, Faye O, Sall AA, Neyts J, Nguyen L, Kaptein SJF, Lambrechts L. 2021. Recent African strains of Zika virus display higher transmissibility and fetal pathogenicity than Asian strains. Nat Commun 12:916.

25. Jagger BW, Miner JJ, Cao B, Arora N, Smith AM, Kovacs A, Mysorekar IU, Coyne CB, Diamond MS. 2017. Gestational Stage and IFN-λ Signaling Regulate ZIKV Infection In Utero. Cell Host Microbe 22:366–376.e3.

26. Szaba FM, Tighe M, Kummer LW, Lanzer KG, Ward JM, Lanthier P, Kim IJ, Kuki A, Blackman MA, Thomas SJ, Lin JS. 2018. Zika virus infection in immunocompetent pregnant mice causes fetal damage and placental pathology in the absence of fetal infection. PLoS Pathog 14:e1006994.

27. Sones JL, Davisson RL. 2016. Preeclampsia, of mice and women. Physiol Genomics 48:565–572.

28. Casazza RL, Lazear HM, Miner JJ. 2020. Protective and Pathogenic Effects of Interferon Signaling During Pregnancy. Viral Immunol 33:3–11.

29. Coan PM, Ferguson-Smith AC, Burton GJ. 2005. Ultrastructural changes in the interhaemal membrane and junctional zone of the murine chorioallantoic placenta across gestation. J Anat 207:783–796.

30. Rossant J, Cross JC. 2001. Placental development: lessons from mouse mutants. Nat Rev Genet 2:538–548.

31. Wells AI, Coyne CB. 2018. Type III Interferons in Antiviral Defenses at Barrier Surfaces. Trends Immunol 39:848–858.

32. Bedford T. 2021. Real-time tracking of Zika virus evolution: https://nextstrain.org/zika.

33. van der Linden V, Pessoa A, Dobyns W, Barkovich AJ, Júnior HV, Filho EL, Ribeiro EM, Leal MC, Coimbra PP, Aragão MF, Verçosa I, Ventura C, Ramos RC, Cruz DD, Cordeiro MT, Mota VM, Dott M, Hillard C, Moore CA. 2016. Description of 13 Infants Born During October 2015-January 2016 With Congenital Zika Virus Infection Without Microcephaly at Birth - Brazil. MMWR Morb Mortal Wkly Rep 65:1343–1348.

34. Hirsch AJ, Roberts VHJ, Grigsby PL, Haese N, Schabel MC, Wang X, Lo JO, Liu Z, Kroenke CD, Smith JL, Kelleher M, Broeckel R, Kreklywich CN, Parkins CJ, Denton M, Smith P, DeFilippis V, Messer W, Nelson JA, Hennebold JD, Grafe M, Colgin L, Lewis A, Ducore R, Swanson T, Legasse AW, Axthelm MK, MacAllister R, Moses AV, Morgan TK, Frias AE, Streblow DN. 2018. Zika virus infection in pregnant rhesus macaques causes placental dysfunction and immunopathology. Nat Commun 9:263.

35. Walker CL, Little ME, Roby JA, Armistead B, Gale M, Rajagopal L, Nelson BR, Ehinger N, Mason B, Nayeri U, Curry CL, Adams Waldorf KM. 2019. Zika virus and the nonmicrocephalic fetus: why we should still worry. Am J Obstet Gynecol 220:45–56.

36. Kwon JY, Aldo P, You Y, Ding J, Racicot K, Dong X, Murphy J, Glukshtad G, Silasi M, Peng J, Wen L, Abrahams VM, Romero R, Mor G. 2018. Relevance of placental type I interferon beta regulation for pregnancy success. Cell Mol Immunol 15:1010–1026.

37. Racicot K, Mor G. 2017. Risks associated with viral infections during pregnancy. J Clin Invest 127:1591–1599.

38. Racicot K, Aldo P, El-Guindy A, Kwon JY, Romero R, Mor G. 2017. Cutting Edge: Fetal/Placental Type I IFN Can Affect Maternal Survival and Fetal Viral Load during Viral Infection. J Immunol 198:3029–3032.

39. Buchrieser J, Degrelle SA, Couderc T, Nevers Q, Disson O, Manet C, Donahue DA, Porrot F, Hillion KH, Perthame E, Arroyo MV, Souquere S, Ruigrok K, Dupressoir A, Heidmann T, Montagutelli X, Fournier T, Lecuit M, Schwartz O. 2019. IFITM proteins inhibit placental syncytiotrophoblast formation and promote fetal demise. Science 365:176–180.

40. Crow YJ. 2011. Type I interferonopathies: a novel set of inborn errors of immunity. Ann N Y Acad Sci 1238:91–98.

41. Ávila-Pérez G, Nogales A, Park JG, Márquez-Jurado S, Iborra FJ, Almazan F, Martínez-Sobrido L. 2019. A natural polymorphism in Zika virus NS2A protein responsible of virulence in mice. Sci Rep 9:19968.

42. Hertzog J, Dias Junior AG, Rigby RE, Donald CL, Mayer A, Sezgin E, Song C, Jin B, Hublitz P, Eggeling C, Kohl A, Rehwinkel J. 2018. Infection with a Brazilian isolate of Zika virus generates RIG-I stimulatory RNA and the viral NS5 protein blocks type I IFN induction and signaling. Eur J Immunol 48:1120–1136.

43. Esser-Nobis K, Aarreberg LD, Roby JA, Fairgrieve MR, Green R, Gale M. 2019. Comparative Analysis of African and Asian Lineage-Derived Zika Virus Strains Reveals Differences in Activation of and Sensitivity to Antiviral Innate Immunity. J Virol 93:e00640–19.

44. Chen J, Liang Y, Yi P, Xu L, Hawkins HK, Rossi SL, Soong L, Cai J, Menon R, Sun J. 2017. Outcomes of Congenital Zika Disease Depend on Timing of Infection and Maternal-Fetal Interferon Action. Cell Rep 21:1588–1599.

45. Aid M, Abbink P, Larocca RA, Boyd M, Nityanandam R, Nanayakkara O, Martinot AJ, Moseley ET, Blass E, Borducchi EN, Chandrashekar A, Brinkman AL, Molloy K, Jetton D, Tartaglia LJ, Liu J, Best K, Perelson AS, De La Barrera RA, Lewis MG, Barouch DH. 2017. Zika Virus Persistence in the Central Nervous System and Lymph Nodes of Rhesus Monkeys. Cell 169:610–620.e14.

46. Diamond MS, Farzan M. 2013. The broad-spectrum antiviral functions of IFIT and IFITM proteins. Nat Rev Immunol 13:46–57.

47. Schoggins JW, MacDuff DA, Imanaka N, Gainey MD, Shrestha B, Eitson JL, Mar KB, Richardson RB, Ratushny AV, Litvak V, Dabelic R, Manicassamy B, Aitchison JD, Aderem A, Elliott RM, García-Sastre A, Racaniello V, Snijder EJ, Yokoyama WM, Diamond MS, Virgin HW, Rice CM. 2014. Pan-viral specificity of IFN-induced genes reveals new roles for cGAS in innate immunity. Nature 505:691–695.

48. Schoggins JW, Wilson SJ, Panis M, Murphy MY, Jones CT, Bieniasz P, Rice CM. 2011. A diverse range of gene products are effectors of the type I interferon antiviral response. Nature 472:481–485.

49. Zhu J, Zhang Y, Ghosh A, Cuevas RA, Forero A, Dhar J, Ibsen MS, Schmid-Burgk JL, Schmidt T, Ganapathiraju MK, Fujita T, Hartmann R, Barik S, Hornung V, Coyne CB, Sarkar SN. 2014. Antiviral activity of human OASL protein is mediated by enhancing signaling of the RIG-I RNA sensor. Immunity 40:936–948.

50. Ivashkiv LB, Donlin LT. 2014. Regulation of type I interferon responses. Nat Rev Immunol 14:36–49.

51. Zani A, Zhang L, McMichael TM, Kenney AD, Chemudupati M, Kwiek JJ, Liu SL, Yount JS. 2019. Interferon-induced transmembrane proteins inhibit cell fusion mediated by trophoblast syncytins. J Biol Chem 294:19844–19851.

52. Hong S, Banchereau R, Maslow BL, Guerra MM, Cardenas J, Baisch J, Branch DW, Porter TF, Sawitzke A, Laskin CA, Buyon JP, Merrill J, Sammaritano LR, Petri M, Gatewood E, Cepika AM, Ohouo M, Obermoser G, Anguiano E, Kim TW, Nulsen J, Nehar-Belaid D, Blankenship D, Turner J, Banchereau J, Salmon JE, Pascual V. 2019. Longitudinal profiling of human blood transcriptome in healthy and lupus pregnancy. J Exp Med 216:1154–1169.

53. Ander SE, Diamond MS, Coyne CB. 2019. Immune responses at the maternal-fetal interface. Sci Immunol 4:eaat6114.

54. Carbaugh DL, Zhou S, Sanders W, Moorman NJ, Swanstrom R, Lazear HM. 2020. Two Genetic Differences between Closely Related Zika Virus Strains Determine Pathogenic Outcome in Mice. J Virol 94:e00618–20.

55. Liu Y, Liu J, Du S, Shan C, Nie K, Zhang R, Li XF, Zhang R, Wang T, Qin CF, Wang P, Shi PY, Cheng G. 2017. Evolutionary enhancement of Zika virus infectivity in Aedes aegypti mosquitoes. Nature 545:482–486.

56. Collette NM, Lao VHI, Weilhammer DR, Zingg B, Cohen SD, Hwang M, Coffey LL, Grady SL, Zemla AT, Borucki MK. 2020. Single Amino Acid Mutations Affect Zika Virus Replication In Vitro and Virulence In Vivo. Viruses 12:E1295.

57. Lemos D, Stuart JB, Louie W, Singapuri A, Ramírez AL, Watanabe J, Usachenko J, Keesler RI, Sanchez-San Martin C, Li T, Martyn C, Oliveira G, Saraf S, Grubaugh ND, Andersen KG, Thissen J, Allen J, Borucki M, Tsetsarkin KA, Pletnev AG, Chiu CY, Van Rompay KKA, Coffey LL. 2020. Two Sides of a Coin: a Zika Virus Mutation Selected in Pregnant Rhesus Macaques Promotes Fetal Infection in Mice but at a Cost of Reduced Fitness in Nonpregnant Macaques and Diminished Transmissibility by Vectors. J Virol 94:e01605–20.

58. Duggal NK, McDonald EM, Weger-Lucarelli J, Hawks SA, Ritter JM, Romo H, Ebel GD, Brault AC. 2019. Mutations present in a low-passage Zika virus isolate result in attenuated pathogenesis in mice. Virology 530:19–26.

59. Kuo L, Jaeger AS, Banker EM, Bialosuknia SM, Mathias N, Payne AF, Kramer LD, Aliota MT, Ciota AT. 2020. Reversion to ancestral Zika virus NS1 residues increases competence of Aedes albopictus. PLoS Pathog 16:e1008951.

60. Grubaugh ND, Ishtiaq F, Setoh YX, Ko AI. 2019. Misperceived Risks of Zika-related Microcephaly in India. Trends Microbiol 27:381–383.

61. Lauck M, Switzer WM, Sibley SD, Hyeroba D, Tumukunde A, Weny G, Taylor B, Shankar A, Ting N, Chapman CA, Friedrich TC, Goldberg TL, O’Connor DH. 2013. Discovery and full genome characterization of two highly divergent simian immunodeficiency viruses infecting black-and-white colobus monkeys (Colobus guereza) in Kibale National Park, Uganda. Retrovirology 10:107.

62. Wang F, Flanagan J, Su N, Wang LC, Bui S, Nielson A, Wu X, Vo HT, Ma XJ, Luo Y. 2012. RNAscope: a novel in situ RNA analysis platform for formalin-fixed, paraffin-embedded tissues. J Mol Diagn 14:22–29.

